# Single-cell T cell receptor sequencing of paired tissue and blood samples reveals clonal expansion of CD8+ effector T cells in patients with calcific aortic valve disease

**DOI:** 10.1101/2023.07.08.548203

**Authors:** Francesca Bartoli-Leonard, Sarvesh Chelvanambi, Tan Pham, Mandy E Turner, Mark C Blaser, Massimo Caputo, Masanori Aikawa, Amanda Pang, Jochen Muehlschlegel, Elena Aikawa

## Abstract

Calcific aortic valve disease (CAVD) is a complex cardiovascular pathology, culminating in aortic stenosis, heart failure and premature mortality, with no comprehensive treatment strategy, except valve replacement. While T cells have been identified within the valve, their contribution to pathogenesis remains unclear. To elucidate the heterogenous phenotype of the immune populations present within patients with CAVD, deep phenotypic screens of paired valve and peripheral blood cells were conducted via flow cytometry (n=20) and immunohistochemistry (n=10). Following identification of a significant population of memory T cells; specifically, CD8+ T cells within the valve, single cell RNA sequencing and paired single T cell receptor sequencing was conducted on a further 4 patients on CD45+ CD3+, CD4+ or CD8+ T cells. Through unsupervised clustering, 7 T cell populations were identified within the blood and 10 identified within the valve. Tissue resident memory (T_RM_) T cells were detected for the first time within the valve, exhibiting a highly cytotoxic, activated, and terminally differentiated phenotype. This pan-pro-inflammatory signal was differentially identified in T cells originating from the valve, and not observed in the blood, indicative of an adaptive, local not-systemic inflammatory signature in CAVD patients. T cell receptor analysis identified hyperexpanded clones within the CD8+ T cell central memory (T_CM_) population, with T_RM_ cells comprising the majority of large and medium clonal expansion within the entire T cell population. Clonal interaction network analysis demonstrated the greatest proportion of clones originating from CD8+ T cell effector memory (T_EM_) and CD4+ naïve / T_CM_ populations and ending in the CD8+ T_RM_ and CD8+ T_CM_ clusters, suggesting a clonal expansion and predicted trajectory of T cells towards a tissue resident, cytotoxic environment within the valve. CDR3 epitope predictive analysis identified 7 potential epitope targets, of which *GALNT4* and *CR1L* have previously been implicated in a cardiovascular context as mediators of inflammation. Taken together, the data identified T cell sub-populations within the context of CAVD and further predicted possible epitopes responsible for the clonal expansion of the valvular T cells, which may be important for propagating inflammation in CAVD.

## Introduction

Calcific aortic valve disease (CAVD) is a complex cardiovascular pathology, resulting in aortic stenosis, subsequent heart failure and premature mortality^1^. Characterized by fibro-calcific leaflet remodeling and reduced left ventricle ejection fraction, CAVD progresses from aortic valve sclerosis to aortic stenosis (AS), with 1 in 4 adults over 65 years old currently affected^2^. While traditional risk factors such as lipid dysregulation, diabetes, smoking and chronic kidney disease all increase the likelihood of developing CAVD^3^, inflammation has been shown to play a critical role in perpetuating the fibrosis and later calcification within the valve, with anti-inflammatory trials such as the CHIANTI [NCT05162742] study highlighting the benefit of reducing inflammation in CAVD^4^. Despite these advancements, surgical intervention is still the first choice of treatment for most, which comes with heightened risk in a multimorbid population^5^. Unlike atherosclerosis, in which lipid lowering treatment has been shown to reduce pathological burden, statins have limited efficacy in delaying the progression of AS and CAVD^4,6^, thus requiring alternative strategies to address this unmet clinical need^7^. Moreover, the question of autoimmunity within the pathogenesis of CAVD still remains to be elucidated. Activated T lymphocytes have been shown to localize with calcific regions within the valve, preceding extracellular matrix remodeling and the deposition of calcium^8,9^. Valvular interstitial cells which line the fibrosa, the side predisposed to calcific lesions, express HLA-DR: a major histocompatibility complex class II protein, which primarily activates CD8+ T cells^10^. Bulk T cell repertoire analysis has previously identified the presence of a clonally expanded activated CD8+ T cell population within the aortic valves^11,12^; however, the phenotype of these leukocytes and their crosstalk with peripheral T cells is unknown. Due to the slow progression of AS and CAVD, questions have dutifully been raised regarding identifying a lymphocyte population able to promote prolonged and consist inflammation in such a manner. Tissue resident memory (T_RM_) T cells (CD8^+^CD103^+^CD69^+^CD49a^+^)^13,14^, phenotypically distinct from effector memory (T_EM_) and central memory T cells (T_CM_), have been shown to produce anti-exhaustion signals, allowing for continued localized inflammation^13^, however they have yet to be identified within the valve. Thus, this study sought to build upon these finding and asses the T cell landscape within the valve leaflet^15^, elucidating the dominant adaptive inflammatory mechanisms that contribute to the progression of CAVD. These findings may aid in the identification of new druggable targets, offering a much needed alternative to surgical intervention.

## Methods

### Study Population

Twenty-four participants who were diagnosed with CAVD undergoing valve replacement surgery were included in this study. Patient demographics are presented in Table 1. Severe aortic stenosis defined as aortic valve area between 0.6–0.8 cm^2^ and mean gradient 40–50 mm Hg. 20 patients were assigned in the flow cytometry arm of the study, with a further 4 assigned to the single cell RNA sequencing (scRNAseq) arm (Figure 1A). Peripheral blood was collected in EDTA vacutainers prior to surgery and single leaflet aortic valve samples were collected and placed in high glucose DMEM (Gibco) and processed immediately. Surgical specimens were collected from two centers, Brigham and Women’s Hospital and Beth Israel Deaconess Medical Center, and approved by respective ethics committees (IRB#2011P001703, IRB#2018-P-000280, respectively). Surgical inclusion criteria included a haematocrit > 25%, and the patient aged between 20-90 years at time of surgery. Exclusion criteria consisted of propr TAVR/AVR, or under the age of 20. All patients provided informed consent for sample collection and data analysis, in accordance with the Declaration of Helsinki.

**Figure 1.**
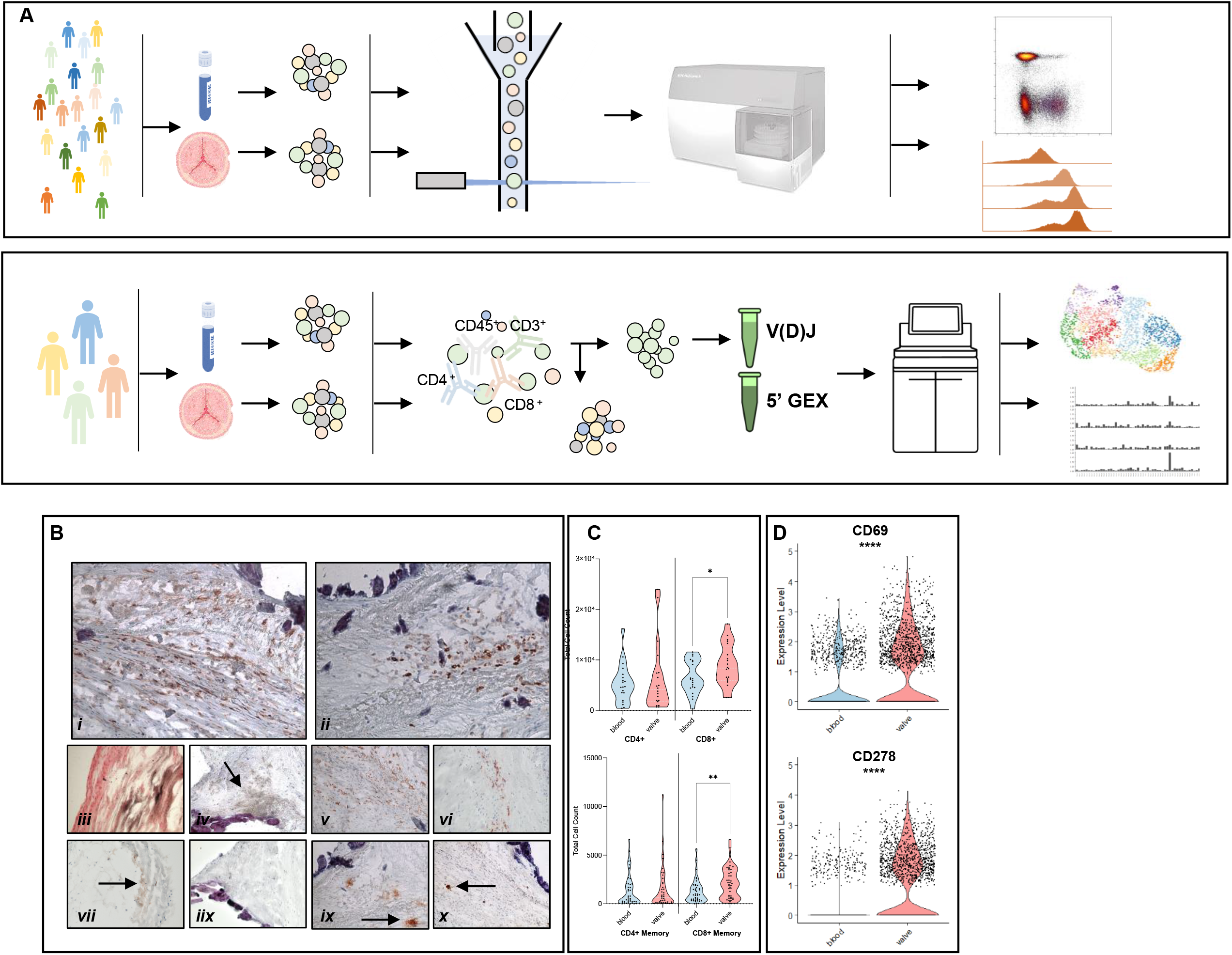
CD8 memory T cell population is present within the blood and increased within the valve. A) Schematic workflow on sample processing and analysis. 20 paired valve and PBMC samples from calcific aortic valve disease patients underwent flow cytometry analysis to phenotype the immune profile. A further 4 donor PBMC and valve leaflet were processed for single cell analysis. Cells were isolated and underwent Fluorescence-Activated Cell Sorting (FACS) to isolate T cells. V(D)J and GEX libraries were then produced and sequenced. 20 donors underwent flow cytometry analysis to phenotype the immune profile. B) Histological staining confirms presence of immune cells (n=10); i) CD4, ii) CD8, iii) Von Kossa, iv) CD20, v) CD68, vi) DC-SIGN, vii) CD123 iix) Siglec-8 ix) Neutrophil elastase x) mast cell tryptase. C) Total cell counts of CD4 and CD8 T cells and memory CD4 and CD8 T cells determined by the presence of CD45RO within the valve and blood (paired T test, n=20). D) Violin plot of single cell gene analysis, CD69 expression on sorted T cells within the valve and blood (Wilcoxon signed-rank test, n=4). Data presented distribution density plots. *p < 0.05, ** p < 0.01. **** p < 0.0001

**Table 1.** Baseline clinical and echocardiographic parameters of tissue donors. 24 donors included in both scRNA and flow cytometry analysis.

|  |  |
| --- | --- |
| **Age, years** | 65 ± 10 |
| **Sex, male (%)** | 18 (75) |
| **Ethnicity, Caucasian, (%)** | 21, (88) |
| **Diabetes, n (%)** | 5 (21) |
| **Hyperlipidmia, n (%)** | 13 (54) |
| **Hypertension, n (%)** | 18 (75) |
| **Ever Smoked, n (%)** | 10 (42) |
| **Previous MI, n (%)** | 1 (5) |
| **Previous PCI, n (%)** | 1 (5) |
| **LVEF (%)** | 60 ± 12 |
| **HbA1c (mmol/mol)** | 5.53 ± 0.68 |
| **Creatinine (mg/dL)** | 0.91 ± 0.37 |
| **Beta-Blocker n (%)** | 9 (38) |
| **Statins n (%)** | 16 (67) |
| **ACE Inhibitor n (%)** | 7 (29) |
| **Angiotensin II receptor Blocker n (%)** | 7 (29) |
| **Bisphosphate n (%)** | 0 (0) |
| **BAV, n (%)** | 14 (58) |
| **Diagnosed CAS, n (%)** | 12 (50) |
| **"Severe" Aortic Stenosis n(%)** | 19 (79) |
| **Peak Ao Transvalvular velocity (m/s)** | 3.68 ± 0.92 |
| **Aortic Valve Area (cm)** | 1.07 ± 1.43 |
| **Ao Peak Gradient (mmHg)** | 61.93 ± 29.95 |
| **Ao Mean gradient (mmHg)** | 43.05 ± 13.01 |
| **Mean/Peak gradient** | 0.60 ± 0.06 |
| **"Severe" aortic incompetence, n (%)** | 10 (42) |
| **Previous AA, n (%)** | 6 (25) |
| **Previous CABG, n (%)** | 6 (25) |
| **Previous Mitral Valve Surgery, n (%)** | 6 (25) |

### Flow Cytometry

Cells were thawed at 37□°C and immediately processed downstream following established protocols^16^. Briefly, cells were counted and suspended at 1×10^6^/mL in sodium-azide free DPBS, with BD Horizon^™^ Fixable Viability Stain (1:1000) at room temperature for 15 minutes protected from light. Cells were then washed in cell stain buffer (Biolegend, USA) and resuspended in 100uL of the same buffer. Samples were then split into 7 FACS tubes and incubated with 100uL of diluted Human TruStain FcX Fc blocking reagent according to manufacturer’s instructions (Biolegend, USA) at 4□°C for 10 minutes protected from light. 100uL of surface antibodies diluted in Brilliant Stain Buffer (BD Biosciences, USA) were added to each vial and incubated at 4□°C for 30 minutes protected from light. Cells were then washed twice in cell stain buffer and for surface-only stain panels, resuspended in 4% paraformaldehyde and incubated at 4□°C for 10 minutes protected from light. Cells were then washed again in sodium-azide free DPBS and resuspended in 300uL of DPBS. Following surface staining and washing, intracellular staining was conducted. Cells were resuspended in 1x Fix/Perm TF Buffer (BD Biosystems, USA) and incubated at 4□°C for 50 minutes protected from light. Cells were then washed in 1x Perm/Wash TF Buffer (BD Biosystems, USA) and incubated with intracellular antibodies, diluted in 1x Perm/Wash TF buffer at 4□°C for 50 minutes protected from light. Cells were then washed twice in cell stain buffer and resuspended in 300uL of DPBS. All samples were pipet filtered prior to acquisition on a CyTek Aurora (Cytekbio, USA). All analysis was conducted using FlowJo (v10.9). Complete list of antibodies used are presented in Table S1.

### Immunohistochemistry

Human tissue specimens were collected from BWH patients diagnosed with CAVD under approval of the Brigham and Woman’s ethics committee (IRB#2011P001703). Specimens were collected immediately after surgery and imbedded in optimal cutting temperature compound (OCT) and snap frozen. Fresh-frozen sections (6 μm) were cut using a cryostat (Leica, Germany) and fixed in 4% paraformaldehyde for 4 minutes. Sections were then washed in PBS for 5 minutes, before blocking in 0.3% H_2_O_2_ in PBS for 20 minutes. Slides were then washed in tap water and placed in PBS for 5 minutes. Slides were then blocked in protein block, serum free (Dako, USA) at room temperature for 1h in a humidified chamber. Following blocking, slides were then incubated with primary antibodies (Supplementary Table 1) diluted in PBS with 5% horse serum and 1% BSA at room temperature for 1 hour in a humidified chamber. Slides were then washed three times in PBS for 5 minutes before diluted secondary antibodies (Supplementary Table 1) were added in PBS with 5% horse serum and 1% BSA at room temperature for 1 hour in a humidified chamber. Slides were then washed three times and incubated with streptavidin-HRP (Dako, USA) at room temperature for 30 minutes before washing a further three times in PBS. Finally, slides were developed in AEC (Dako, USA) and washed in tap water. Slides were then counterstained in Gils haematoxylin for 30 seconds before washing twice in tap water, placed in ammonium water for 15 seconds and then washed in water twice again. Slides were mounted using a water-soluble mounting media and imaged on a Zeiss AxioImager microscope.

### Tissue processing and cell collection

Peripheral blood mononuclear cells (PBMC) were isolated using ACK Lysing Buffer (Thermo Fisher, USA). Fresh blood was diluted 1:10 in ACK lysing buffer and rotated at room temperature for 30 minutes until lysis was complete. Samples were then centrifuged at 400g, 4□°C for 5 minutes and the pellet was washed once in PBS before resuspending and freezing in Bambanker media (Bulldog, USA). Fresh aortic valves were collected in DMEM and cut into approximately 1 mm^3^ pieces and placed in 1mg/mL collagenase from Clostridium histolyticum (Sigma, USA) and gently mixed for 1h. Supernatant was then removed, and the pellet was processed (digest 1). The remaining valve tissue was resuspended in collagenase as before and mixed for a further 3 hours. Following the second digestion, supernatant was again processed as before and frozen (digest 2), ready for downstream processing.

### Single cell sorting and preparation

Fluorescence activated cell sorting (FACS) was utilized to isolate T cells from PBMCs and valve cell isolate. Briefly, valve cells (digest 1 and 2, combined) and PBMCs were defrosted, washed once in cell stain buffer (Biolegend, USA) and resuspended in FACS stain buffer with Zombie UV live/dead stain (1:1000, Biolegend, USA) at 4□°C for 15 minutes protected from light. Cells were then washed in FACS stain buffer and resuspended in diluted Human TruStain FcX Fc blocking reagent according to manufacturer’s instructions (Biolegend, USA) at 4□°C for 10 minutes protected from light. T cells were then stained for CD45 (AF488, Biolegend, USA), CD4 (AF700, Biolegend, USA), CD8 (PE, Biolegend, USA) and CD3 (AF647 Biolegend, USA) and incubated at 4□°C for 30 minutes protected from light. Cells were then washed in stain buffer twice and resuspended in stain buffer and sorted immediately on the FACS Aria II (BD Biosystems, USA) into FACS tubes. Following sorting, CD45+ CD3+ CD8+ and CD45+CD3+CD8 cell populations were combined per donor and immediately processed downstream.

### Single cell RNA-Sequencing

Sorted cells were immediately processed following the Chromium Next GEM Single Cell 5’ Dual Index protocol (10x Genomics, USA). Briefly, gel beads (GEMs) were generated by combining 1100 cells/uL with single cell VDJ 5’ Gel beads, a master mix, and portioning oil onto a Chromium Next GEM kit K via a Chromium Controller (10x Genomics). Isolated paired immune receptor sequences and whole transcriptome for V(D)J and gene expression (GEX) libraries were encapsulated within the GEM. Immediately following GEM generation, gel beads were dissolved, and the co-partitioned cell was lysed via heating samples on a thermocycler (Veriti, Applied Biosystems) at 53□°C 45 minutes, 85□°C for 5 minutes before cooling on ice. Additionally, the master mix released primers containing an illumine TruSeq read I sequence, a 16nt 10x barcode, a 10nt unique molecular identifier (UMI) and a 13nt template switch oligo (TSO). Following this, GEMs are broken, pooled and Silane magnetic beads (SPRI Select, Beckman Coulter) were used to purify the 10X barcoded first-strand cDNA from poly-adenylated mRNA and DNA from cell surface protein specificity feature barcode from the post GEM-RT reaction mixture. cDNA is then amplified to generate sufficient material for both TCR and GEX libraries, which DNA is amplified for cell surface protein feature barcode libraries. Amplified full-length cDNA from poly-adenylated mRNA is used to enrich full-length V(D)J barcoded segments via PCR amplification with *primers* specific to TCR constant regions. Enzymatic fragments and size selection was then used to generate variable length fragments that collectively span the V(D)J segments with P5, P7, i5 and i7 sample indexes and an illumina R2 sequence via end repair, A-tailing, adaptor ligation and sample index PCR, with the final libraries containing P5 and P7 priming sites used in illumina sequencing. Amplified full-length cDNA from poly-adenylated mRNA is used to generate 5’ gene expression library. Enzymatic fragmentation and size selection are used to optimize the cDNA amplicon size prior to GEX library construction. P5, P7, i5 and i7 sample indexes and an illumina R2 sequence are added via end repair, A-tailing, adaptor ligation and sample index PCR, with the final libraries containing P5 and P7 priming sites used in Illumina sequencing. Following completion of all Illumina ready dual index libraries, all libraries were assessed by KAPA quantification (Roche) and sequenced on an Illumina NovaSeq S4, samples were duel indexed with a 10bp index length, 1% Phi-X spike in, with 2000 cells per sample recovery target.

### Single-cell RNA-Seq data processing

Reads were mapped to the GRCH38 human genome using the Cellranger toolkit (10X genomics, version 7.01) with the default parameters. Dead cells and other artefacts were filtered out as previously described^17^ Quality control was performed downstream in the Seurat pipeline^18^ implemented in R 4.1.3 environment R Studio. Briefly, the number of genes, UMIs, and the proportion of mitochondrial genes for each cell was calculated. Cells with a low number of covered genes (<500), or high mitochondrial counts (mt-genes >5%) were excluded from downstream analysis, the matrix was then normalized via log-transformation and scaled in Seurat using the SCT method. Gene dispersion was calculated and cells with a high dispersion and coverage (2000 genes) were included in the principal component analysis (PCA). Cells lacking T cell markers CD45, CD3, CD8 or CD4 were excluded from downstream analysis. Seurat objects for GEX were produced per patient, before merging for blood or valve. Merged objects were normalized using the SCT method and integrated using RPCA reduction and clustered according to the Seurat vignette. Dimensional reduction was visualized via uniform manifold approximation and project (UMAP) reduction. VDJ sequencing data was merged with the Seurat objects via the combineExpression function of scRepetoire^19^. Valve leaflets, blood, and the merged valve-blood dataset was phenotyped using the celldex^20^, human atlas and blueprint encode packages^21,22^. Concurrently, phenotypes were also predicted by the iCEllR^23^ and SingleR^20^ packages, and predicted phenotypes were then crossed referenced with a manually curated dataset of key phenotypic genes taken from landmark papers within the single cell field^24–34^. Clonotyping analysis was conducted via scRepetoire, visualized via circulize^35^ and ggplot2. Briefly, contigs from the TCRA and TCRB chains were combined to create a single list object with the TCR genes; comprised of the VDJC genes and CDR3 sequences by cell barcode. Clonotypes were quantified per relative and absolute abundance per patient and sample type, with the dynamic clonotype movement between PBMC T cells and valve determined through alluvial mapping. Clonal space homeostasis, determined by the relative space occupied by the clones at specific proportions. The ClonalNetwork function was utilized to interrogate the network interaction of clonotypes shared between clusters along the single cell dimensional reduction, with the function showing the relative proportion of clones that come from starting nodes, shown by the arrow direction. Clonal networks were filtered by proportion, showing only those considered *medium* clonally expanded and above^38^. Movement of clonotypes between clusters was produced by chordDiagram() function in *circulize viz* and shown per donor. Enriched interaction pathway analysis; single cell gene set enrichment analysis (GSEA) was completed using Escape^36^. Briefly, molecular signature databases were downloaded from GSEABase geneSet Collection objects and the molecular signature database^37^. Visualization was produced using dittoPlot^38^.

### Definition of clonal expansion

Clonotype expansion was determined as per default scRepitoire settings. Following the combining of the TCR and CDR3 barcode to produce a TCR signature, clonotypes within blood, valve and merged Seurat objects were categorized as follows: Single defined as a unique clonotype, small defined as 1 < x ≤ 5 clonotypes; medium defined as 5 < x ≤ 20 clonotypes; large defined as 20 < x ≤ 100 clonotypes; and hyperexpanded determined as x > 100 clonotypes.

### Epitope prediction modelling

Epitope binding prediction was conducted via TCRmatch through the IEDB interface. Briefly, TCRMatch compares CDR3b sequences against a curated CDR3b sequence library within the IEDB to predict shared epitope specificity. Matches are determined by sequence similarity and scored via a comprehensive k-mer comparison through the VDJ server.

### Data availability

Raw scTCR-seq dataset from the patients are not publicly available to protect research participant privacy/consent. Please contact the corresponding author for enquires.

## Results

Recent single cell analysis studies in human atherosclerotic plaques have demonstrated an accumulation of T cells within diseased regions compared to non-pathological vessels^39^, however little is known regarding the T cell population in late stage aortic stenosis, subsequent calcification and their contribution to prolonged inflammation within the valve. Histological analysis (n=10) of classical immune cell markers were assessed in diseased valve to confirm the presence of inflammatory infiltrates (Fig 1B). CD4 and CD8 staining were present in areas consistent with the most damaged parts of the valve, adjacent to the highest-grade calcification; shown via von Kossa staining and structural degeneration. CD20 B cell staining was minimal, with few positive cells observed. Flow cytometry analysis conducted on 20 paired blood/valve donors identified 4 subtypes of B cells: memory, naïve, transitional, and plasmablast (Fig S1). CD27+ IgD^lo^ memory B cells presented as less than 5% of the total live cells in both blood and valve. IgD+ CD27- naïve B cells were significantly (p<0.05) more abundant in valves compared to the blood, as were CD38+ CD24- plasmablasts (p<0.05). Transitional B cells represented the greatest population of B cells within both the valve and blood and were significantly greater within the valve (p>0.05), however variation was high between donors. As previously demonstrated via IHC, the CD68+ monocyte/macrophage fraction was highly prevalent within the valve leaflets. Flow cytometry analysis of whole blood and valve demonstrated a significant increase in CD192^+^ HLA-DR^lo^ classic monocytes in the blood compared to the valve (p>0.01) (Fig S1). Intermediate monocytes, characterized via CD192+ HLA-DR+ accounted for a significantly greater population; 14% of the live cells within the valve, compared to 8% within the blood (p>0.05). Finally, nonclassical CD192lo, CD36+ monocytes were significantly reduced in the valve compared to the blood (p>0.001). Staining with DC-SIGN evidenced the presence of dendritic cells within the valve though to a much lower count than monocytes (25% vs >1%). All subtypes of DC combined represented less than 1% of live cells within either the blood or valve, with no significant difference observed between them. Notably however, when compared as ratios, CD141+ CD208+ cDC1 mature DCs were observed in 4x greater prevalence than immature CD141+ CD208- cDC1 DCs (p>0.01) and similarly compared to CD208+ CD1c+ cDC1 mature DCs (p>0.05). CD123+ basophils, Siglec-8 eosinophils, neutrophil elastase-stained neutrophils, and mast cells stained via mast cell tryptase were infrequently observed within the tissue (Fig 1B). Flow cytometry analysis of granulocytes identified very few; <1% of live cells as CD123+ CD11b+ basophils, CD11b+ Siglec-8+ eosinophils within either the valve or blood. Conversely, the CD11b+ CD16+ neutrophil population within the blood with significantly greater than that in the valve (p>0.0001) and represented >96% of granulocytes within the blood, whereas within the valve basophil, neutrophil, and eosinophil were similarly distributed. T cell populations, characterized by CD45+ CD3+ CD8+ or CD4+ were observed within both the blood and valve via flow cytometry, with CD8+ T cells significantly more abundant within the valve compared to the blood (p<0.05) (Fig 1C). More specifically, CD8+ memory T cells expressing the CD45RO marker were significantly higher in the valve compared to the blood (p>0.01).

Following the identification of a divergent population of T cells within the valve compared to the blood, it was important to understand the T cell population and if they were passively present or active within the valve leaflets. CD69 is a marker of a possible recent antigen encounter and activation; known to rapidly elevate after active engagement of the TCR with activation signals^38^. Gene expression of CD69, detected via scRNAseq, was significantly increased (p<0.0001) in valve compared to blood T cells. Similarly, CD278, recently demonstrated to be critical in the activation and homing of T cells^38^, was significantly increased within valve T cells compared to the blood. These results taken in conjunction with the prevalence of T cells within the valve suggests that T cells are present and actively engaging in TCR-specific interactions within the diseased valve.

To further investigate the population of T cells and elucidate their phenotypic distribution, 13-plex flow cytometry was conducted on whole blood and valve tissue digests (n=20). Treg populations, defined by their CD4+ CD23+ FoxP3+ markers, represented less than 1% of the live CD45+ valve cells and were significantly more abundant in the blood (p>0.01) (Fig 2A). Markers of T cell activation were assessed in both CD4+ and CD8+ T cell subsets. Activated CD4+ HLA-DR+ T cells and CD45RO+ CD197- effector memory cells were at significantly greater abundance in the valve compared to the blood (p<0.001, p<0.05), conversely both CD45RO+ CD197+ central memory cells were at greater abundance in the blood compared to the valve (p>0.001) (Fig 2B). Naïve CD197+ CD45RO-CD4+ T cells were significantly greater in the blood than the valve (p>0.01). Activated CD4+ T cells represent over a quarter of CD4+ T cells within the valve, compared to < 5% within the blood. CD8+ T cells were also characterized within the blood and valve (Fig 2B). Activated, effector and naïve CD8+ T cells presented no significant total count difference between the blood and valve; however, proportionally, activated and effector cells represented a greater percentage of CD8+ T cells than they did in the blood. Central memory T cells were significantly greater abundance within the valve compared to the blood (p>0.05), and conversely, effector memory CD8+ T cells had 3x the prevalence in the blood than the valve (p>0.001). Similarly, when compared as a percentage to the whole CD8+ subtype, effector memory CD8+ T cells represented close to 40% of CD8+ T cells within the blood, whereas this value is less than 10% within the valve. Central memory CD8+ T cells were relatively 12% more expanded within the valve than the blood.

**Figure 2.**
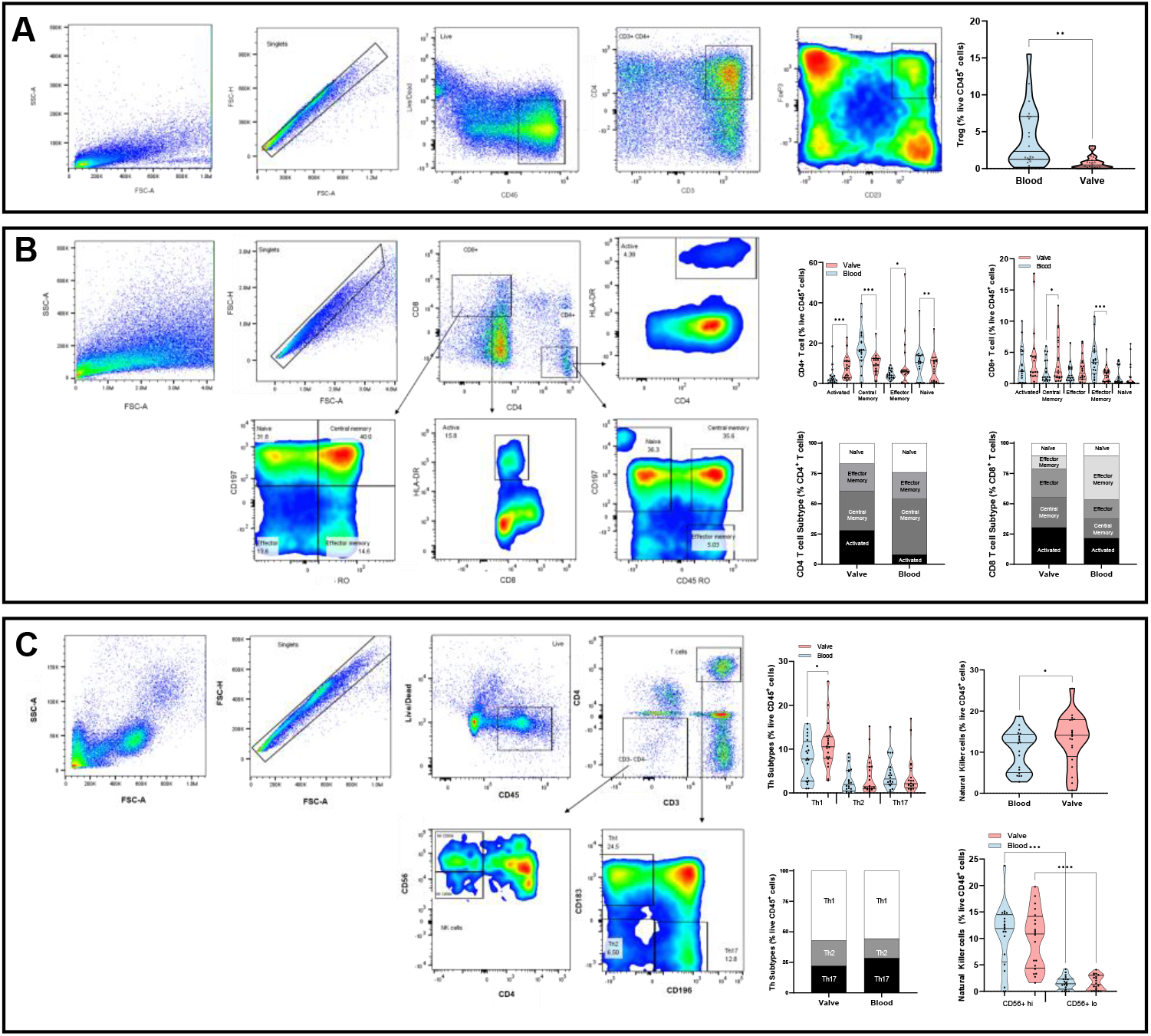
Flow cytometry analysis identifies distinct T cell populations within the valve and peripheral blood. A) Treg cell gating strategy identified by CD45+CD3+CD4+CD25+FoxP3+ phenotyping. B) Gating strategy for activation of CD4 and CD8 T cells. CD45RO+CD197+ determined central memory, CD45RO+CD197- determined effector memory, presence of HLA-DR determined activated T cells and CD45RO-CD197+ determined naïve T cells. C) Th1 subtype was determined by the presence of CD183 and CD196 on CD4 T cells. Natural Killer (NK) cells were determined by the presence of CD56 and categorized as CD56hi and CD56lo. Data presented distribution density plots with interquartile range or as ratios. Ratios calculated based on live CD45+ cell count. Statistical analyses were performed using paired-T tests corrected for multiple comparisons (n=20). *p < 0.05, ** p < 0.01 ***p < 0.001.

CD4+ subtypes can further be broken down to helper subtypes: Th1 which predominately mediates cellular immune response; Th2 which potentiates humoral response; and Th17 which is a plastic subtype characterized by its production of IL-17. CD4+ T helper cell subtype counts were similar between the valve and the blood, with more Th1 cells present within the valve than the blood (p<0.01). Notably, the ratio between Th1, Th2, and Th17 remained similar within both, with Th1 cells representing over 50% of the helper cell subtype (Fig 2C). Finally, innate lymphocytic natural killer (NK) cells were more prevalent within the valve than the blood (p>0.01), with CD56hi+ NK cells representing the majority of NK cells in both the valve and the blood (p>0.001, p>0.001).

Given the expansion of memory cells specific to the valve, a further four paired blood-valve donors were assessed to single cell resolution to assess their clonal expansion and transcriptomic signature. Aortic valve leaflet and peripheral blood were digested and filtered to a single cell solution, before sorting to elucidate a pure CD45+CD3+ CD4+ / CD8+ population via FACS (Fig S2). Single cell analysis confidently identified 2637 cells and 2554 cells within the blood and valve, respectively. Uniform manifold approximation and projection (UMAP) was used to display the data (Fig 3A, D). Unsupervised clustering was conducted via Seurat, with 7 T cell clusters identified within the blood, and 10 within the valve. The relative proportions of T cells within each cluster is represented in Fig 3B and 3E.

**Figure 3.**
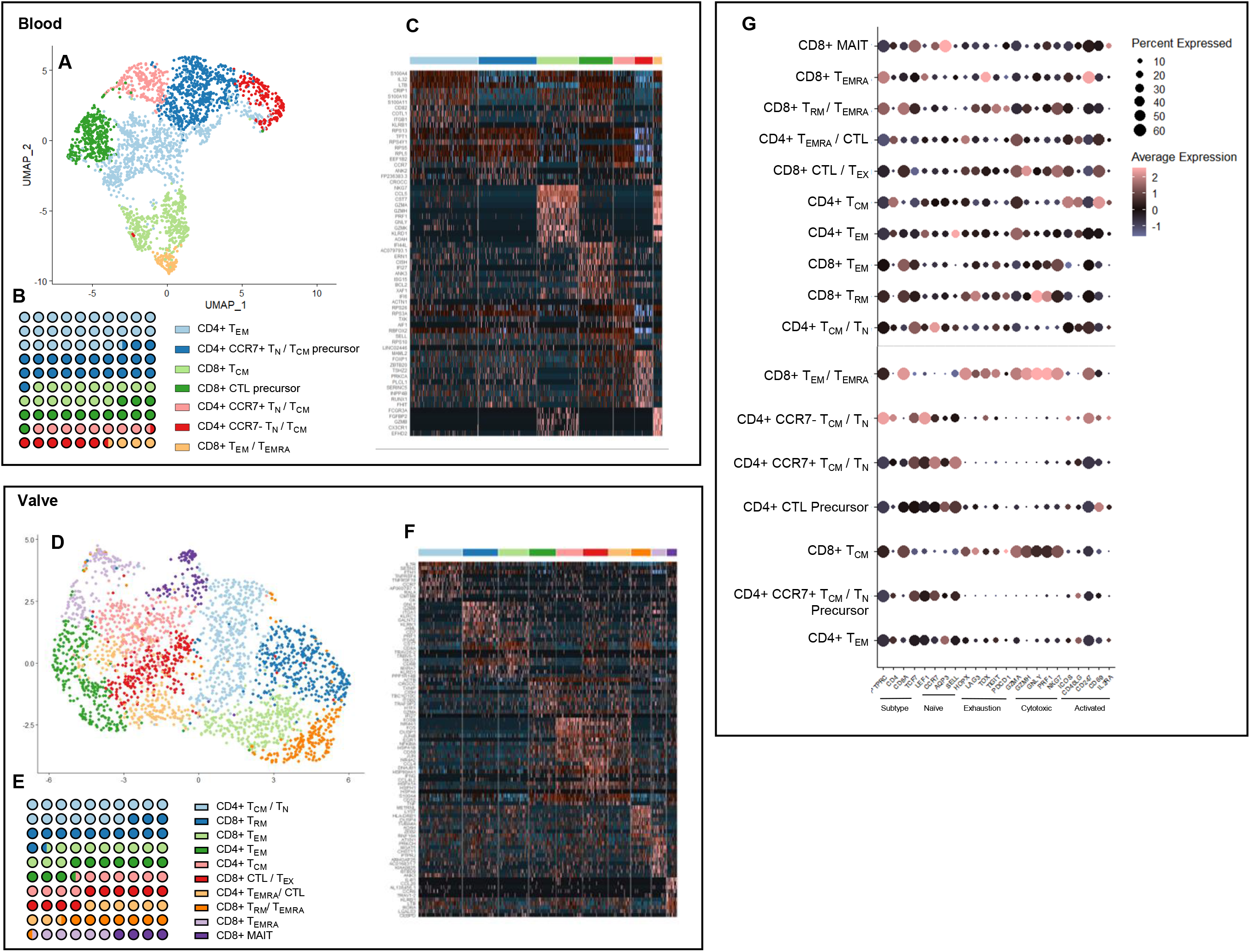
Single cell sequencing of T cells identifies distinct populations between valve and blood. A, D) UMAP depicting distinct T cell clusters with the peripheral blood and valve tissue resulting from unsupervised clustering. B, E) Proportional representation of the cell counts within the clusters C, F) Heap map with average expression of top 10 most differentiability expressed gene for each cluster. G) Dot plot average expression of T cell function associated genes, split by cluster (blood: n=2637, 4 donors, valve: n=2554, 4 donors). Data presented as means unless stated otherwise.

Within the blood, cells were firstly identified through their CD4 / CD8 expression (Fig S3, A, D) with further in-depth phenotyping conducted through *celldex*, *iCellR*, *SingleR* and a manually curated dataset. Within the blood, three naïve, three memory, and one cytotoxic population were identified (Fig 3A). Differential gene expression of blood CD4+ T_EM_ T cells had key effector memory markers *ITGB1, IGALS1* and *S100A4* ^40,41^ and exhibited a loss of naïve *CCL5* and *LEF1*. Moreover, effector markers *IL-32, S100A11*, regulatory *TNFRSF4* and classical *CXCR3* were increased in the T_EM_ cluster, representing 27.5% of the T cell population. CD4+ T_CM_ / T_N_ precursor cells, which accounted for 23.5% of the T cell population within the blood, expressed leukocyte adhesion/tracking marker CCR7 and naïve marker *LEF1*, but not *CCL5*, suggesting a movement away from complete nativity. Notably, cytotoxic activation markers *GZMA, GZMH* and *GZMK* were significantly decreased in this cluster, with minimal classical T_CM_ marker *IL-7R*. CD4+ T_N_ / T_CM_ clusters were stratified via *CCR7* expression, with *CCR7*+ T_N_ / T_CM_ sharing 79% similarity with *CCR7*- T_N_ / T_CM_, collectively representing 15% of the T cell population. Both populations exhibited high levels of classical naïve cell markers *SELL* and *LEF1*, but with a relative loss of *CCL5*. Neither population exhibited cytotoxic gene expression; *GMZA*, *GMZB* or *GNLY*, with significantly lower expression of inflammatory interferon markers compared to other clusters. Late-stage activation marker *HLA-DR* was decreased in both clusters, with early activation marker *CD69* and the gamma chain of *CD25* demonstrated limited expression, suggesting a move towards activation. By comparison, 14% of T cells were clustered into a CD8+ T_CM_ population and exhibited high levels of cytotoxic markers; *GZMH, GZMA, GNLY* and *GZMK*. Naïve markers *CCR7* and *LEF1* were lost in T_CM_ cluster, with cells expressing lymphocyte-attracting chemokine *CCL4* and CCL5, indicative of activated central memory cells. CD8+ CTL precursor cells highly expressed interferon IFI27 and *IFI44L* and *ISG15*, known to promote antigen specific CTL interferon activation, but lacked active CTL *PRF1*, *GZMA* and *GZMB* markers. Finally, 3.5% of the T cell population, which clustered as CD8+ T_EM_ / T_EMRA,_ expressed classical markers of terminal differentiation, including *CCR7*-, *CD28*-, *CD27*- and highly expressed cytotoxic markers: *PRF1* and *NKG7* (Fig 3C).

The majority of the populations within the valve tissue were memory T cells, with one innate-like T cell population identified (Fig 3D). CD4+ T_CM_ and CD4+ T_CM_ /T_N_ share a 40% homology, representing 10% and 17% of the T cell population, respectively. T_CM_ /T_N_ expressed nativity markers *CCR7*+ and moderate *SELL*+, which are lacking in the T_CM_ cluster. Conversely, memory markers *CCL4* and *CD69* are significantly greater in the T_CM_ then T_CM_ /T_N_ cluster, however neither expresses a strong cytotoxic phenotype, indicative of specific antigen activation (Fig 3F). CD8+ T_RM_ cells represent 14.5% of the T cell population and express the core transcriptional residency program of T_RM_ cells, characterized by high expression of adhesion and retention markers (*CXCR6*, *CD49a*, *CD103, JAML*), infiltration (*KLRC1*) and cytotoxicity (*GZMB*, PRF1), as well as a decrease of tissue egress markers (*CCR7*, *SELL*, *S1PR1*). Similarly, CD8+ T_RM_/ T_EMRA_ cells demonstrated increased adhesion, retention, and cytotoxicity markers, as well as a loss of *CCR7*, *CD28*, *CD27*, suggesting terminal differentiation and representing 8% of the T cell population. A smaller cluster of 5.5% of the T cell population were identified as CD8+ T_EMRA_, presenting similar markers to the previous cluster without the presence of *CD49a, CD69, CD103* and a decreased expression of terminal homing marker *CX*CR6. CD4+ T_EMRA_ /CTL exhibited a reduction in *CD28* and *CCR7* and concurrently an increase in proinflammatory gene activation markers *FOS*, *TNFα* and *IL-32*, as well as classical CTL marker *GZMA*. 10% of the CD8+ T cell population exhibited both cytotoxic markers (*GMZA, IFNG, NKG7*), exhaustion marker (*KLRG1*), and cell cycle inhibition markers (*NR4A2* and *NR4A3*) alongside high levels of *CCL4*, all of which when taken together is indicative of a CTL phenotype^42^. Both CD4+ and CD8+ T_EM_ cells demonstrated a loss of *CCR7* and an increase in rest marker *CCL5*. CD8+ T_EM_ cells exhibited a more cytotoxic phenotype with greater expression of *PRF1, HOPX, KLRD1, EOMES*, and *NKG7*, coupled with a loss of naivety marker *SELL*. Similarly, CD4+ T_EM_ exhibited an increase in memory markers *LFA-1, CD52*, and *HLA-DR,* coupled with less cytotoxic markers with the exception of *IFNG*. Finally, mucosal-associated invariant (MAIT) T cells represented 4% of the total T cells within the valve, and expressed classical MAIT markers (*KLRB1, CCR6* and *CXCR6*) with a lack of *CCR7* and *CD62L*, and the invariant *TRAV1-2* TCR chain.

Identified clusters were further categorized into T cell states (Fig 3G). As expected, precursor and naïve populations within the blood expressed the greatest level of naïve cell markers (*LEF1*, *ACP3*, *CCR7*, *SELL*, *HOPX*). Conversely, cytotoxic (*GZMA, GZMH, GNLY, PRF1, NKG7*) and exhaustion (*LAG3, TOX, TIGIT, PDCD1*) markers were highly associated with the valvular memory cells and the CD8+ populations, with little to no expression seen in the CD4+ or CTL precursor population. Activation marker *CD247* was highly expressed in all clusters with T cells isolated from the valve exhibiting greater expression of costimulatory markers *ICOS, CD40L* and *CD60*, indicative of an immunological active population.

Markers of maturation from naïve to terminal (progressing from *CCR7, SELL, CD27, CXCR4, CXCR3* to *CXCR6*) were assessed in the blood and valve tissue T cells (Fig. 4A). Notably, blood T cells exhibited greater naïve marker expression, *CCR7* and *SELL*, compared to the valve (p>0.0001, p<0.0001). Activation of T cells via the TCR/CD3 receptor induces expression of *CD27*, which decreases after prolonged activation. *CD27* was expressed both within the valve and blood but significantly greater within the blood (p<0.0001). Expression on *CXCR4* and *CXCR3*, both markers of terminal differentiation and maturation, were higher in valve tissue than blood (p>0.0001, p<0.0001). Similarly, when examining these markers within each cluster, activation of cytokine and proliferative markers (Fig S3, B, E) was observed more frequently within the valve than the blood. CD69 activation was observed in the majority of valve cell clusters (p>0.001) and notably absent in blood clusters, with the exception of the CD8+ CTL precursor (p>0.01) (Fig S3, C, F).

**Figure 4.**
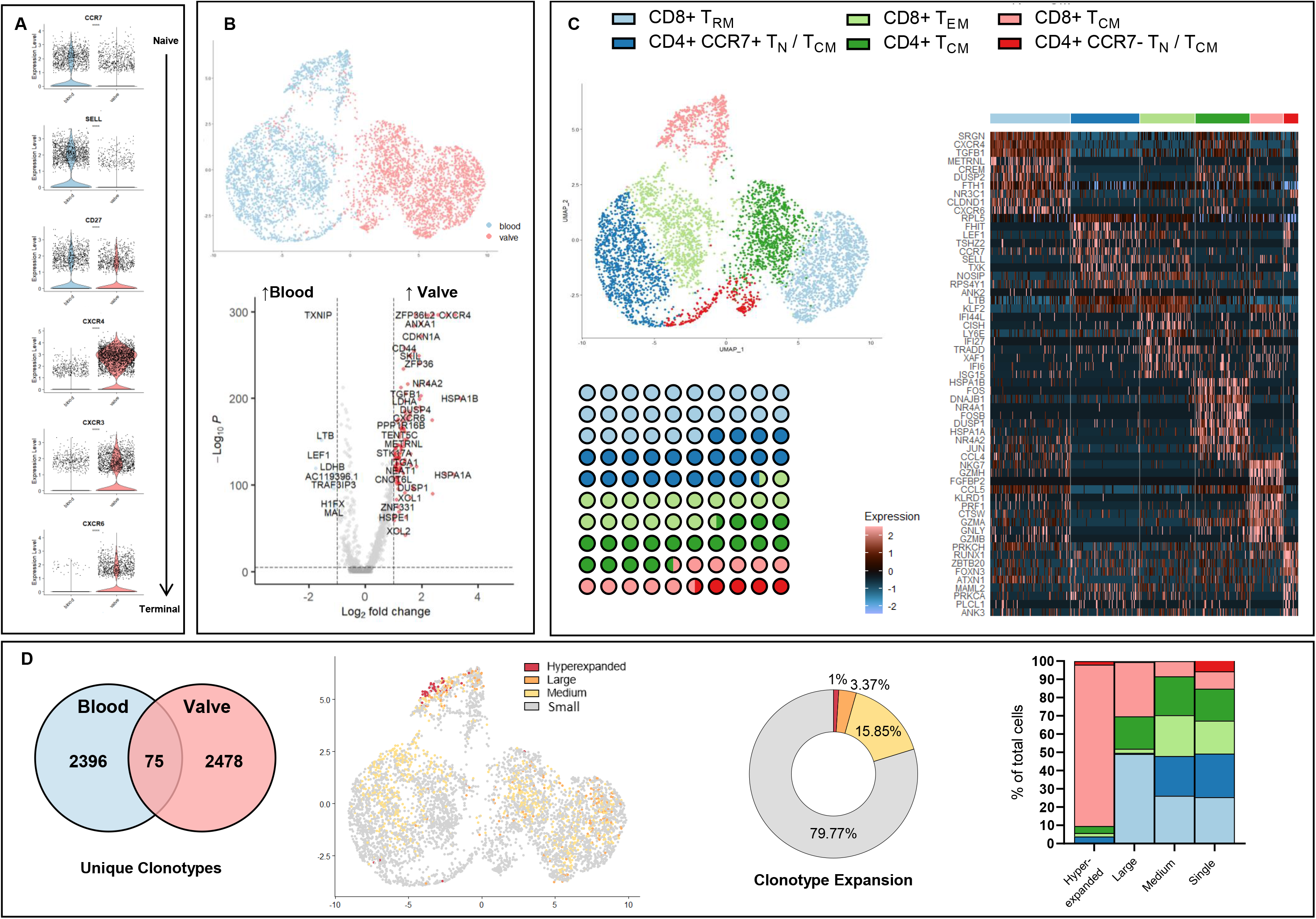
Clonal expansion occurs predominantly within the CD8+ tissue resident memory and central memory T cells. A) Violin plots of differentiation markers; CCL7, SELL, CD27, CXCR4, CXCR3 and CXCR6 within the blood and valve. B) UMAP of blood and valve T cells. Volcano plot with differentially expressed genes in valve or blood T cells. C) UMAP visualization of unsupervised clustering revealed six distinct T cell populations within the merged blood–valve T cell population. Proportional representation of the cell counts within the clusters shown below. Heap map with average expression of top 10 most differentially expressed genes for each cluster. D) Total number of unique clonotypes identified in blood and valve T cells, with 75 unique clonotypes shared between both. Expansion of clonotypes mapped onto UMAP with proportional representation of percentage of clonotype expansion within all cells. Clonal expansion mapped to cell type defined by unsupervised UMAP as a percentage of all cells within the expansion bracket. Data presented as means unless stated otherwise. Statistical analyses were performed using paired-T tests. **** p < 0.0001. Clonotype expansion levels: Single; one occurrence, medium (>0.1% to ≤ 1%), large (>1% and ≤10%), hyperexpanded (>10%). (blood: n=2637, 4 donors, valve: n=2554, 4 donors).

Examination of blood and valve T cells together demonstrated little transcriptomic overlap between the two source tissues (Fig 4B) with notable differential expression of inflammatory markers (*CXCR4, GNLY, GZMA*) in the valve compared to the blood. Unsupervised clustering of the merged dataset identified six CD45+ CD3+ T cell clusters (Fig 4C, S3G), complied to assess clonal expansion between the peripheral and local T cell populations. Similar to the analysis of only blood T cells, CD4+ clusters were characterized as *CCR7*+/- naïve and central memory phenotypes, expressing classical naïve markers *SELL*, *LEF1* and *CCR7*, which represented a combined 44% of the T cells identified. CD8+ T cells were all identified as memory cells, with the T_RM_ population the largest CD8+ cluster (26%). Similar to the analysis of only valve T cells, in the merged dataset T_RM_ cells had increase expression of tissue homing *CXCR6*, *CD103* and *CD49a* markers as well as proinflammatory *PRF1* and *GNLY*. Notably CD8+ T_CM_ expressed a strong inflammatory phenotype (*PRF1, GZMA, GNLY*) and trafficking marker *KLF2*. CD8+ T_EM_ were notably less cytotoxic, with *TNF*α and interferon associated genes notably increased in comparison to other clusters. Cytokine and proliferative activation were observed more strongly in populations originating from the valve than the blood, in line with the single tissue analysis (Fig S3 H, I).

Single cell sequencing of the TCR was achieved concurrently with gene expression analysis. 94% and 99% of TCRs on the blood and valve cells were sequenced. Of the 4949 identified clonotypes, 3% (75) were shared between valve and blood (Fig 4D). Clonal expansion of the TCR repertoire was determined through the expansion of both the TCR alpha and beta clonotypes, defined as small, medium, large, and hyperexpanded for those that constituted >0.001, >0.01, >0.1 and 1% of the analyzed T cell population, respectively. Hyperexpanded cells exhibited significantly greater homeostatic space within the blood than the valve, demonstrating a reduction in T cell clonal diversity (*x*^2^=12.348, p=0.000259) (Fig S4A, G) with 21% and 27% of clonal expanded cells present in the valve and blood respective populations compared to 79% and 73% unique diverse clonotypes. Phenotypically, 88% of hyperexpanded clonal population was identified as CD8+ T_CM_ cells. Within the large and medium clonal space repertoires, T_RM_ cells represented the largest proportion of these clonal populations; 49% and 26%, respectively. Notably, CD4+ CCR7- T_CM_ / T_N_ cells exhibited little clonal expansion. Similar results were observed within the blood and valve analysis separately (Fig S4B, H). Clonotype analysis was diverse between donors (Fig S4 C, I); however, contribution of clonotypes within the tissue types was similar (Fig S4 D, J, S5 A). The flow of clonotypes from blood to valve was assessed within each donor, with the top 10 most abundant clonotypes presented (Fig 5A). As expected, donors were heterogenous in clonal diversity, with Donor 3 and 4 having unique top 10 clonotypes. Donors 1 and 2 shared 6 of the top 10 most abundant clonotypes, with the largest proportion of clonotypes in both donors being CILGGGNQFYF. In the combined dataset of all donors, CILGGGNQFYF was again the highest proportionally represented clonotype, and three of the top ten clonotypes in the blood was not expressed within the valve at all, indicating a localized clonal expansion triggered by an epitope specific to the valve. Bidirectional movement of the clonotype distribution via chord diagram is depicted in Fig 5B and S4F, L. CD4+ T_CM_ / T_N_ clusters showed little clonal movement. Conversely, CD8+ T_RM_ and CD8+ T_CM_ exhibited greater interconnection of clusters through their TCR. By overlaying of the clonal interaction network with the UMAP, the directionality of network interactions between different clusters can be visualized (Fig 5C, S4 E, K). CD8+ T_CM_ demonstrated an efflux of the proportion of clones from the starting nodes in this cluster out to CD8+ T_EM_ and CD8+ T_RM_ clusters, with 100 and 150 unique clones moving from the starting note to the ending node. Similarly within the CD8+ T_RM_ population, clonal expansion proportionality moved from both the CD8+ T_CM_ and CD4+ CCR7- T_CM_ / T_N_ populations inwards towards the T_RM_ population, with a small (>50 unique clones) but high proportion (>1%) of clones starting from CD4+ CCR7- T_CM_ / T_N_ and moving into the T_RM_ clusters. Complete clonal movement from individual clusters (Fig S5C, D) was calculated as a delta of overall net movement, with CD8+ T_RM_ and CD4+ CCR7+ T_CM_ / T_N_ having the most an efflux of unique clonotypes transitioning in from other populations, 20 and 18, respectively.

**Figure 5.**
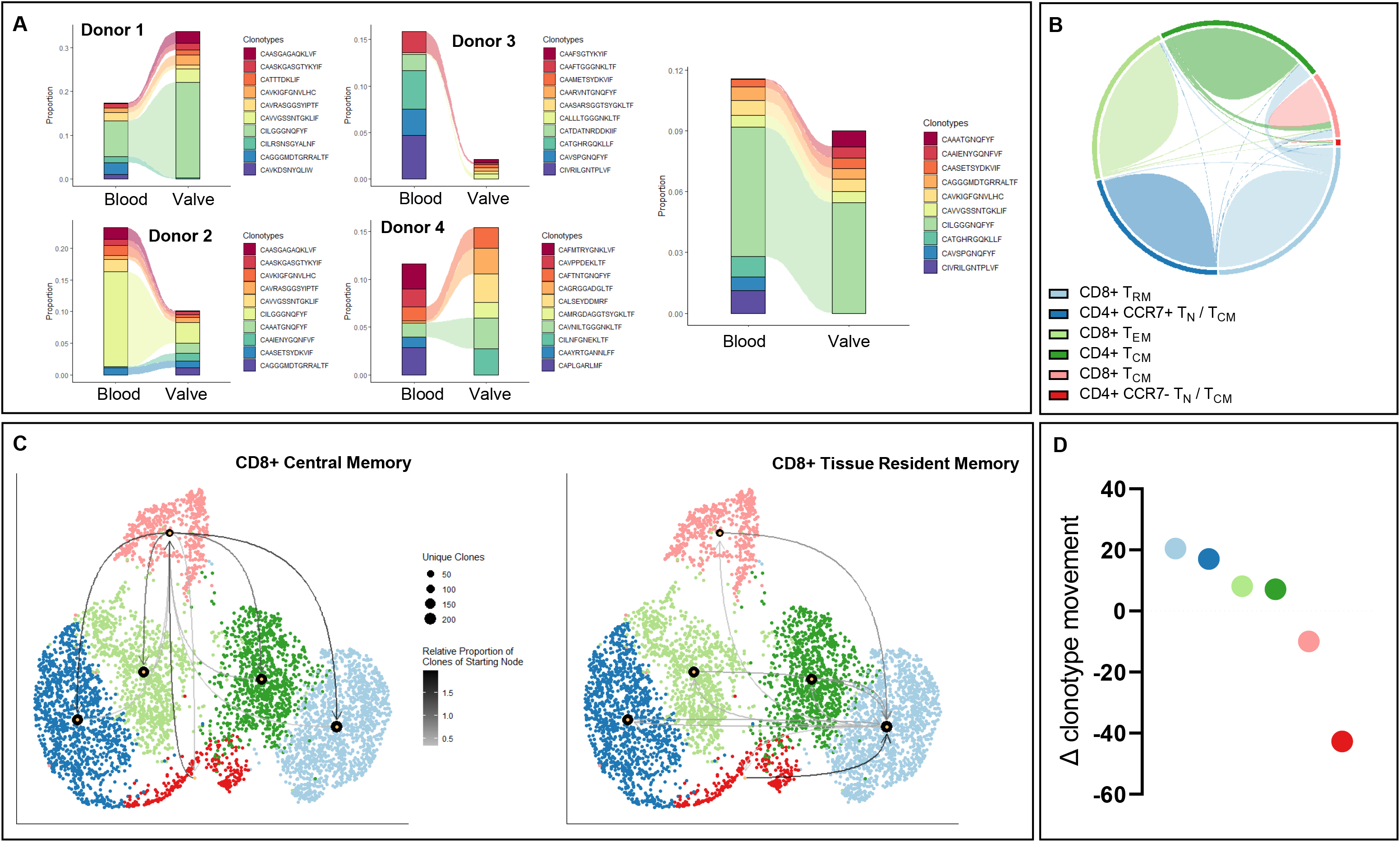
Movement of clonal expansion demonstrates a migration from CD8+ central and effector memory to CD8+ tissue resident memory. A) Dynamic changes between blood and valve T cell clonotypes per patient and combined valve-blood representing the top 10 clonotypes. B) Interconnection of clusters via chord diagram. C) UMAP visualization overlay identifying the network interaction of clonotypes shared between clusters along the single cell dimension reduction. Relative proportion of clones that come from a starting node and end in a different cluster, CD8+ central memory T cell and CD8+ tissue resident memory T cell clonal network visualized. D) Delta of the clonal network calculated. Positive delta demonstrates a movement of clones from other clusters into the named cluster. (blood: n=2637, 4 donors, valve: n=2554, 4 donors).

As the data suggests a consortium of inflammatory T cell populations present within the CAVD, GSEA was conducted to further understand the heterogenous populations present within the valve leaflet tissue (Fig 6A). As expected, CCR7^+/-^ naïve / central memory CD4+ T cells demonstrated a limited inflammatory profile and conversely both CD4+ and CD8+ effector and central memory exhibited a higher level of proinflammatory, interferon-driven pathways. Notably, CD8+ T_RM_ cells were associated with proinflammatory, hypoxic, and metabolic pathways, seen within the Hallmark and Molecular Signature databases^36^. Of specific interest, cytolytic, proinflammatory and reactive oxygen species pathways were increased in CD8+ T_RM_, T_EM_ and T_CM_ clusters, as well as CD4+ T_CM,_ and were characteristically lower in the naïve cell clusters (Fig 6B, p>0.001 for all). Separating clusters by origin identified a greater inflammatory profile within CD8+ T_EM_ originating from the valve tissue, with a greater population of naïve cells originating from the blood identified as less inflammatory than their corresponding valve population. T_RM_ cells were identified solely in the valve tissue, with significant levels of proinflammatory and inflammatory response GSEA pathways enriched (p>0.001, all).

**Figure 6.**
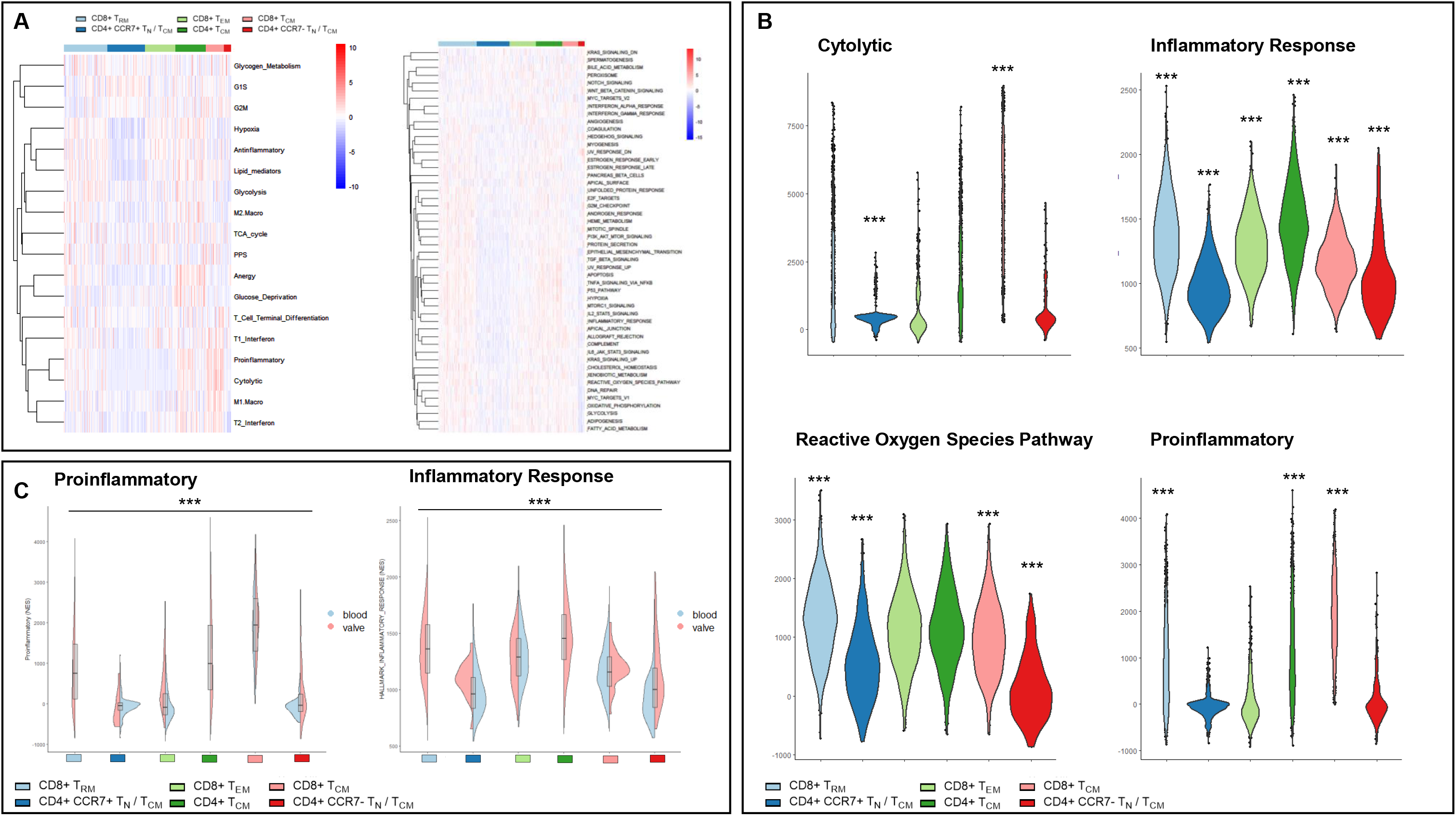
Enriched interaction pathways between T cell clusters identifies inflammatory responses within the valve. A) Heatmap analysis of pathways activated within T cell clusters through GSEA. B) Subset of inflammatory pathways visualized via violin plot. C) Comparison of proinflammatory and inflammatory response GSEA subset by tissue of origin, significance denotes difference between blood and valve enrichment. Significance marked if group significantly different to all other clusters only. (blood: n=2637, 4 donors, valve: n=2554, 4 donors). Statistical analyses were performed using ANOVA. ***p < 0.001.

Predictive analysis of the potential epitope binding partner for CDR3 of the TCRs identified by scRNAseq was conducted via TCRMatch (Table 2). A total of 19 potential epitopes were predicted, corresponding to 7 source molecules. Of the 7 molecules, polypeptide N-acetylgalactosaminyltransferase 4 (*GALNT4*) and complement component receptor 1-like protein (*CR1L*), predicted to be recognized via CILGGGNQFYF and CATGHRGQKLLF, respectively, have previously been identified in relation to T cell activation and subsequent inflammation. *CR1L* plays a role in the regulation of complement dependent cytotoxicity within a cardiac ischemia reperfusion injury model. Fragments of the CR1 was identified as an injury specific neoepitope ^43^, with inhibition suggested to be a possible target for reducing ischemia in injured cardiac systems. Secondly, *GALNT4* has previously been identified in genome wide association studies within to coronary artery disease in relation to decreased platelet and endothelial function^44–46^. Notably, neither source molecules have been identified within the context of CAVD, however given the strong associations between CAD, atherosclerosis and CAVD further investigation into these molecules is warranted.

**Table 2.**
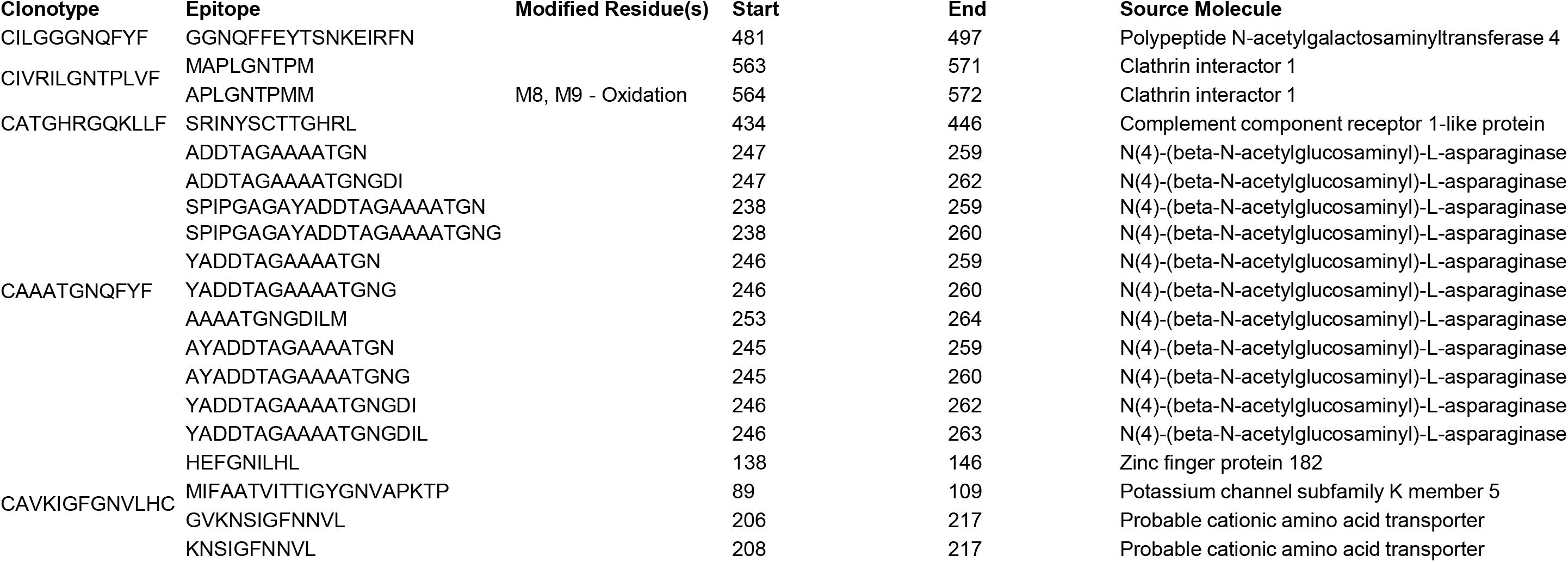
Top 5 unique CDR3 mapped TCRs to known epitopes through TCRMatch.

## Discussion

Current treatments for aortic valve stenosis comprise surgical intervention alone, and has yet to leverage the molecular and mechanistic insight gained from translational research. While CAVD is considered, in part, an inflammatory disease^47^, little research has been conducted to examine the specific immune cell populations at play within the pathogenesis of CAVD, nor as a potential target for pharmacological intervention. Landmark studies^11,48^ in CAVD identified circulating clonally expanded activated and effector memory T cells supporting the relationship between immunogenic events in the pathophysiology of CAVD. However, studies have stopped short of elucidating the transition of T cells and their receptors and their crosstalk between the valve and the blood, and how these populations expand and traverse the environment.

Thus, this study sought to investigate the T cell population within the patient with CAVD from both the systemic circulation and localized within the diseased valvular tissue, in order to better understand whether CAVD is driven by a localized expansion of T cells within the valve, or a migration of clonally expanded T cells from the blood to the valve. Nearly 20% of the T cells detected within the total sequenced cells were identified as clonally expanded, with the majority of single clones originating from naïve populations within the blood. scRNA analysis demonstrated the accumulation of clonally expanded T cells predominantly within those isolated from the blood as opposed to the valve leaflet, which, through predictive network analysis were then identified as T_RM_ and T_CM_ CD8+ subpopulations. The population of T_N_ cells were clustered into both CCR7- and CCR7+ phenotypes. CCR7, a hallmark of naïve cells and a receptor for the constitutive chemokine *CCL19* and *CCL21*, enables migration of T cells to lymphoid organs for antigen presentation via DCs^49^. A lack of CCR7 in T cells has been demonstrated to be indicative of a low adaptive immune response and an abnormal migratory pattern, coupled with increased cell survival following overactivation of T cell in response to a stimuli^50^. The novel population of T_N_ / T_CM_ CCR7- T cells observed in this cohort may be indicative of an exhausted response from the T cell population, promoting a low level but consistent inflammatory environment in which CAVD can progress. T_CM_, both CD4+ and CD8+, were identified within the CAVD population. Both originating from the blood, CD8+ T_CM_ were the majority of the 1% hyperexpanded T cells within the valve, suggesting a preference for pro-inflammatory clonal expansion over naive populations. In contrast to recent studies on atherosclerosis^39^, the results here demonstrate a clonal expansion skewed to CD8+ rather than CD4+ T cells, though the authors did note due to a lack of paired blood-tissue T cells, they could not confidently assess the CD8+ expansion with atherosclerotic plaques^51^. By sub-setting T cells within the context of CAVD, the heterogeneity of T cells within the valve is apparent, with both pro-and anti-inflammatory T cells seeking to regulate or perhaps dysregulate the immune landscape of CAVD.

This study is the first to demonstrate the presence of CD8+ T_RM_ in CAVD, defined by their phenotype, transcriptional profile, and enrichment networks. Notably, these cells display low levels of naïve characteristics, with greater levels of activation (*CD247*, *CD25),* cytotoxicity (*GZMH*, *GNLY*), and residency (*CD103, CD69, CD49a*) markers when compared to other CD8+ T cell subtypes. The CD8+ T_RM_ identified here demonstrate a similar transcriptomic profile to CD8+ T_RM_ cells those found in rejected transplanted allografts^52^, suggesting a similar mechanistic role in low grade inflammatory pathogenesis. Moreover, selective differentiation of naïve T cells to T_RM_ within the valve may be promoted by the presence of transforming growth factor beta (TGFβ) n the aortic valve leaflets^53^, which has been previously shown to promote T_RM_ differentiation within tissues^54^. These findings extend the concept of T_RM_ cells functioning in cases of chronic rejection, to a case of lower grade inflammatory-mediated structural degradation, in which epitopes present on the valvular tissue stimulate perpetually activated T cells. This chronic milieu may contribute to CAVD pathogenesis and progression. Investigation into the potential epitope binding partners of T cells assessed, and T_RM_ cells specifically, identified CILGGGNQFYF as the most frequent clonally expanded CDR3 within the population, which is predicted to bind with *GALNT4*. *GALNT4;* present in the later stages of atherosclerosis is colocalized to atherosclerotic plaques and upregulated in activated monocyte populations. Furthermore, *GLANT4* in the context of atherosclerosis has been identified as a key initiator of mucin-type O-glycosylation, which plays a causal role in the increased susceptibility of pan-cardiovascular disease, concurrently upregulating with the progression of atherosclerosis. However, it is worth noting that all predicted epitopes suggested here are autologous to humans, with no external bacterial or viral epitopes, in contrast to epitope prediction in rheumatic heart valve disease T cell analysis^55^ which has a known viral cause, which adds weight to the suggestion of CAVD being an auto-immune driven pathogenesis. Prediction of epitope TCR binding partners may allow future research to focus on the specific epitope-TCR reaction, giving rise to more targeted therapies in the context of CAVD, an avenue which is yet to be explored.

In summary, this study mapped the T cell repertoire in CAVD, utilizing T cells isolated from paired valve-blood donors, demonstrating that T cells are clonally expanded within the pathology and predicting a directionality of peripheral to local T cell movement targeting specific epitopes. A subset of T cells within the valve expressed hallmarks of T_RM_ cells, expressing specificity to proteins identified within cardiovascular disease. This may mediate an attack on self that triggers the development and progression of CAVD thus suggesting a mechanistic autoimmune role of T cells. Moreover, the data presented here shows that different T cell subsets display contrasting signature characteristics within the valve tissue and the circulation, supporting the role T cells within CAVD and suggesting that T cells may promote both pro-and anti-inflammatory effects within the aortic valve.

### Sources of Funding

This work was supported by the NIH grants R01HL147095, R01HL141917, and R01HL136431 to EA and the Elizabeth Blackwell Institute, University of Bristol, and the Wellcome Trust Institutional Strategic Support Fund (ISSF3, 204813/Z/16/Z) to FBL. BHF Translational Award Grant TA/F/21/210028 and British Heart Foundation Professor of Congenital Cardiac Surgery chair (CH/17/1/32804) to MC.

#### Disclosures

The authors state no disclosures.

## Acknowledgments

N/A.

## Notes

### Competing Interest Statement

The authors have declared no competing interest.

