## Supplemental figures & tables for "Single-cell T cell receptor sequencing of paired tissue and blood samples reveals clonal expansion of CD8+ effector T cells in patients with calcific aortic valve disease"

| Antibodies | Source | Cat # | Fluorophore | Dilution | Assay |
|---|---|---|---|---|---|
| CD4 | Biolegend | 317401 | | 1:100 | IHC |
| CD8 | Biolegend | 372902 | | 1:100 | IHC |
| CD20 | Biolegend | 302301 | | 1:100 | IHC |
| CD68 | Abcam | ab213363 | | 1:1000 | IHC |
| CD123 | Biolegend | 306002 | | 1:100 | IHC |
| DCSIGN | Abcam | ab245115 | | 1:100 | IHC |
| FoxP3 | Biolegend | 320101 | | 1:150 | IHC |
| Siglec-8 | Biolegend | 347202 | | 1:100 | IHC |
| Mast Cell Trypase | Abcam | ab2378 | | 1:1000 | IHC |
| Neutrophil Elastase | Abcam | ab2541 | | 1:150 | IHC |
| Live/dead | BD Biosciences | 565388 | *FVS780* | 5uL per test | FC |
| CD45 | BD Biosciences | 555482 | *FITC* | 5uL per test | FC |
| CD14 | BD Biosciences | 563079 | *BV510* | 5uL per test | FC |
| CD16 | BD Biosciences | 562874 | *BV421* | 5uL per test | FC |
| CD11c | BD Biosciences | 561356 | *PE-Cy7* | 5uL per test | FC |
| CCR2 | BD Biosciences | 747847 | *BB700* | 5uL per test | FC |
| CD36 | BD Biosciences | 555455 | *PE* | 5uL per test | FC |
| CCR6 | BD Biosciences | 551773 | *PE* | 5uL per test | FC |
| CXCR3 | BD Biosciences | 566532 | *BB700* | 5uL per test | FC |
| CD56 | BD Biosciences | 564058 | *BV786* | 5uL per test | FC |
| CD25 | BD Biosciences | 555432 | *PE* | 5uL per test | FC |
| FoxP3 | BD Biosciences | 561182 | *V450* | 5uL per test | FC |
| CD33 | BD Biosciences | 742217 | *BB700* | 5uL per test | FC |
| CD141 | BD Biosciences | 565321 | *BV421* | 5uL per test | FC |
| CD1c | BD Biosciences | 742750 | *BV786* | 5uL per test | FC |
| DC LAMP | BD Biosciences | 558126 | *PE* | 5uL per test | FC |
| CD19 | BD Biosciences | 562947 | *BV510* | 5uL per test | FC |
| CD20 | BD Biosciences | 561175 | *PE-Cy7* | 5uL per test | FC |
| IgD | BD Biosciences | 566538 | *BB700* | 5uL per test | FC |
| CD27 | BD Biosciences | 563327 | *BV786* | 5uL per test | FC |
| CD38 | BD Biosciences | 659478 | *BV421* | 5uL per test | FC |
| CD24 | BD Biosciences | 555428 | *PE* | 5uL per test | FC |
| CD3 | BD Biosciences | 564713 | *BV510* | 5uL per test | FC |
| CD8 | BD Biosciences | 555366 | *FITC* | 5uL per test | FC |
| CD4 | BD Biosciences | 557852 | *PE-Cy7* | 5uL per test | FC |
| CD45RO | BD Biosciences | 560607 | *PerCP-Cy5.5* | 5uL per test | FC |
| CCR7 | BD Biosciences | 566741 | *PE* | 5uL per test | FC |
| CD11b | BD Biosciences | 555388 | *PE* | 5uL per test | FC |
| CD123 | BD Biosciences | 566482 | *BB700* | 5uL per test | FC |
| CD16 | BD Biosciences | 562874 | *BV421* | 5uL per test | FC |
| Siglec-8 | BD Biosciences | 747873 | *BV786* | 5uL per test | FC |
| HLA-DR | BD Biosciences | 564041 | *BV786* | 5uL per test | FC |
| HLA-DR | BD Biosciences | 560651 | *PE-Cy7* | 5uL per test | FC |
| CD3 | Biolegend | 300322100 | *AF647* | 5uL per test | FC / FACS |
| CD8 | Biolegend | 344706 | *PE* | 5uL per test | FC / FACS |
| CD4 | Biolegend | 344622 | *AF700* | 5uL per test | FC / FACS |
| CD45 | Biolegend | 344622 | *AF700* | 5uL per test | FC / FACS |
| Zombie UV | Biolegend | 423108 | *UV 355* | 1:1000 | FC / FACS |

### Supplemental Table 1. Antibody Table

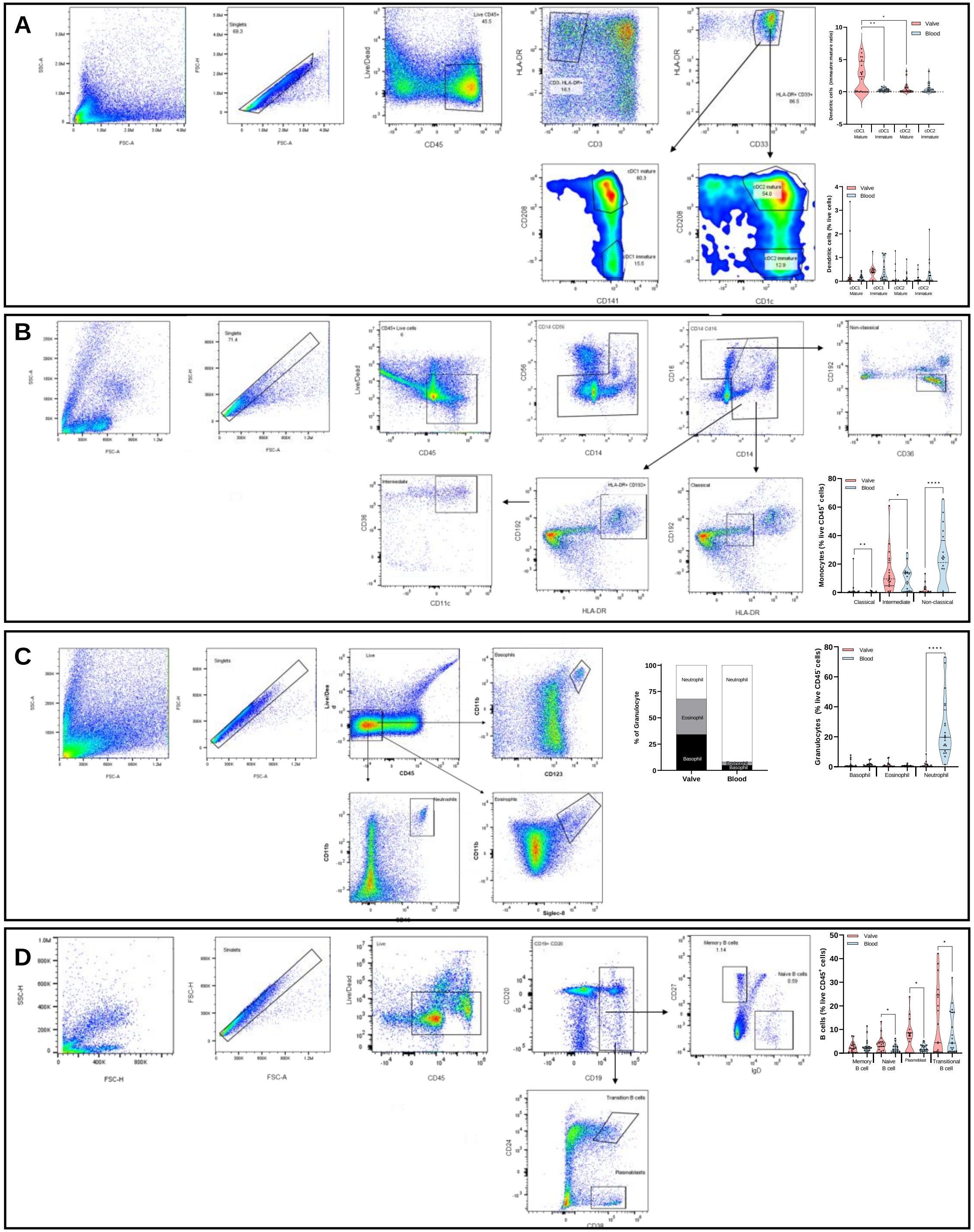


**Supplemental Figure 1. Flow cytometry analysis identifies distinct immune populations within the valve and peripheral blood.**

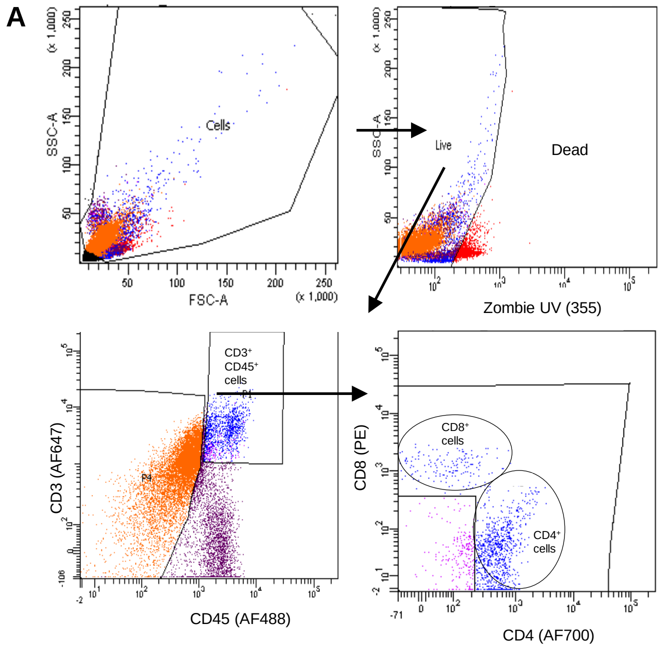


**B**

| Population | Valve # Events | Valve % Live | Blood # Events | Blood % Live |
|---|---|---|---|---|
| All events | 50000 | | 45000 | |
| Cells | 38287 | | 14394 | |
| Live | 19422 | 100 | 13083 | 100 |
| $CD3^+$ $CD45^+$ cells | 1294 | 7 | 3230 | 25 |
| $CD4^+$ & $CD8^+$ cells | 821 | 4 | 2893 | 22 |

**Supplemental Figure 2. Gating strategy for T cell sorting for single cell RNA sequencing populations.**

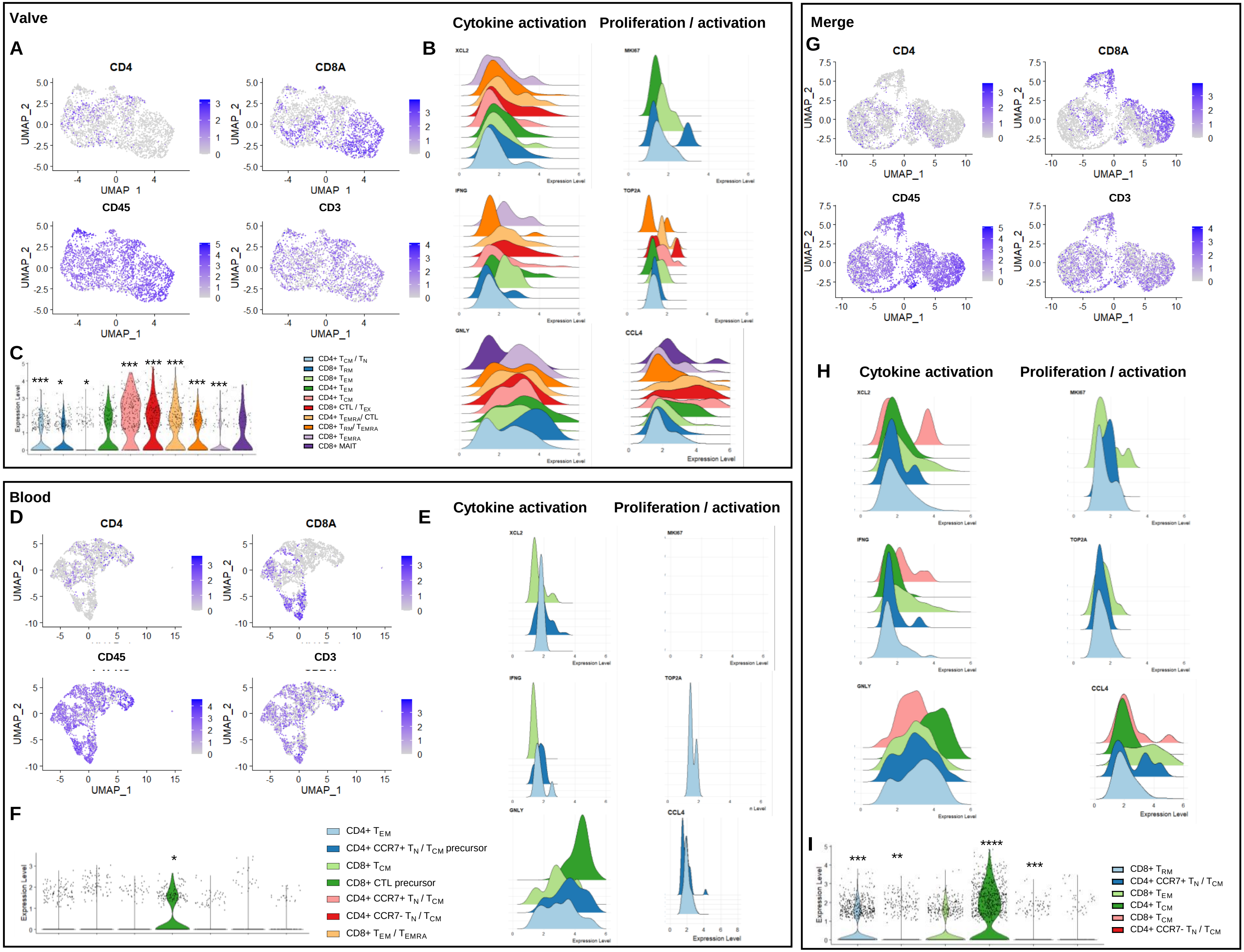


**Supplemental Figure 3. T cell marker distribution and population per cluster.**

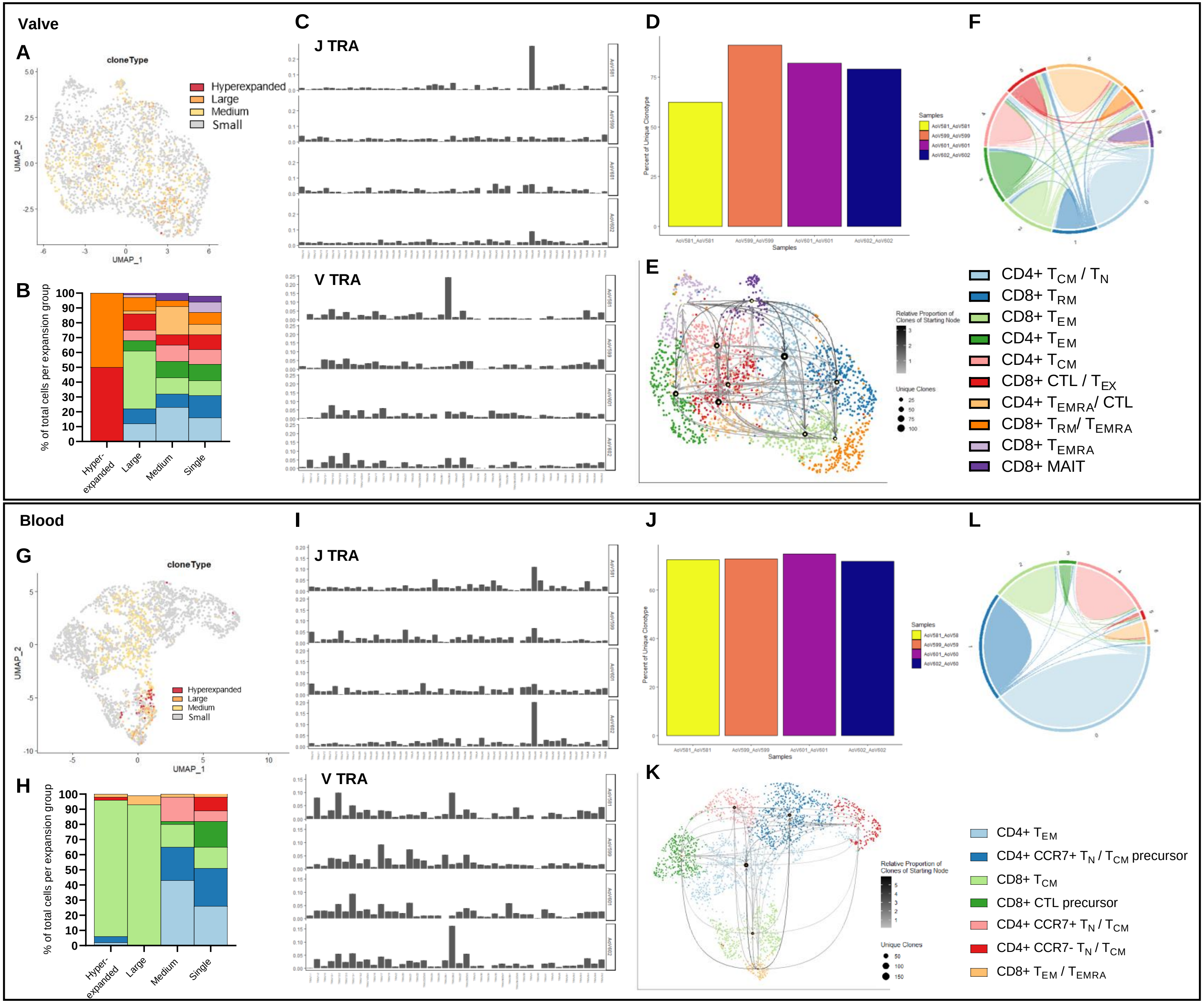


**Supplemental Figure 4. Independent valve and blood clustering identifies clonal expansion and shared clonotype mapping.**

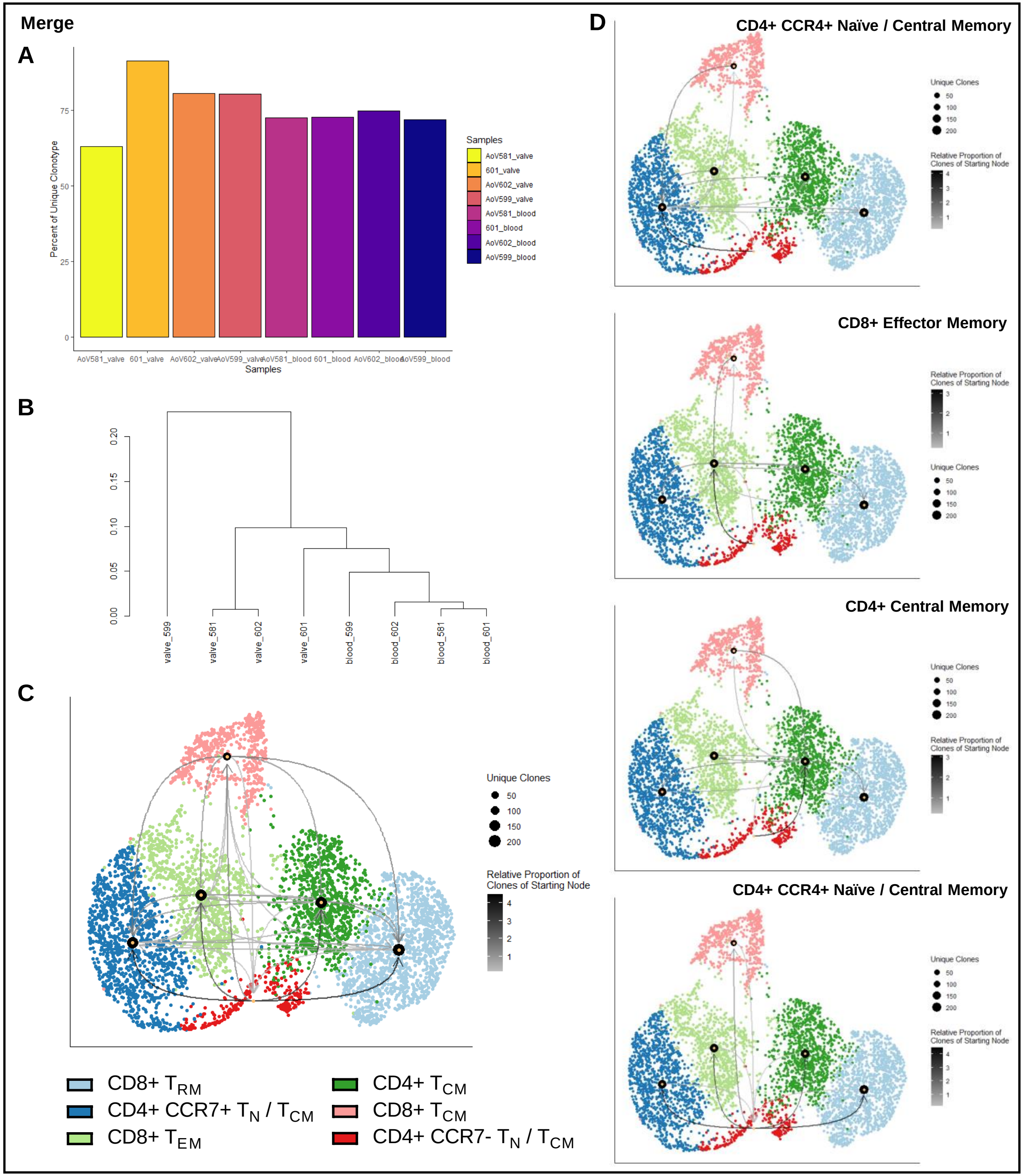


**Supplemental Figure 5. Contribution and distribution of donor samples and shared clonotype mapping between clusters.**
